# *Bradyrhizobium hardenbergiae* sp. nov., isolated from *Hardenbergia violacea* in Australia, represents a novel basal lineage of the *B. elkanii* supergroup

**DOI:** 10.1101/2024.08.26.609660

**Authors:** Bénédicte Lafay, Elina Coquery, Philippe M. Oger

## Abstract

Bradyrhizobia are widespread across the Australian continent, where they are essential to Australian ecosystems by helping legumes to compensate nutrient deficiencies and low fertility of Australian soils. Among the *Bradyrhizobium* genospecies identified during a survey of Australian native rhizobia communities in 1994-1995, genospecies L appeared to be only distantly related to any *Bradyrhizobium* lineages known at the time. We take advantage of the recent sequencing of the genome of strain BDV5419, the original strain corresponding to *Bradyrhizobium* genospecies L, to re-assess this lineage taxonomic status. We characterized further strain BDV5419 based on morpho-physiological traits and determined its phylogenetic relationships with the type strains of the 88 currently known *Bradyrhizobium* species based on sequence comparisons of SSU rRNA genes and complete genomes. The digital DNA–DNA hybridization relatedness with any type strain was less than 33% and both SSU rRNA gene and genome phylogenies confirmed that this strain does not belong to any formerly described species within the *Bradyrhizobium* genus. Whereas its position within the lineage encompassing the *B. elkanii* and *B. jicamae* supergroups is unresolved in the SSU rDNA phylogeny, strain BDV5419 appears to be one of most basal lineages of the *B. elkanii* supergroup in the genome comparison. All data thus support the description of the novel species *Bradyrhizobium hardenbergiae* sp. nov. which type strain is BDV5419^T^ (= CFBP 9111^T^ = LMG 32897^T^), isolated from a nodule of *Hardenbergia violaceae* in Black Mountain Nature Reserve, in Canberra, ACT, Australia.

---

*Bradyrhizobium* bacteria (*Hyphomicrobiales*: *Nitrobacteraceae*) are the most ubiquitous and abundant bacteria in soils (1). Whilst they may live as saprophytes (2), they are foremost known as nodule-inducing nitrogen-fixing bacteria capable of associating symbiotically with Fabaceae (3). They are the most frequently isolated rhizobia (4) and nodulate the widest range of legume genera (5), among which the most basal lineages within the Caesalpinoideae (6). In addition, *Bradyrhizobium* is the only genus found in the wild in symbiotic association with *Parasponia* (Cannabaceae), the sole currently known non-leguminous plants that can establish a nitrogen fixing rhizobial mutualism (7,8). *Bradyrhizobium* is also remarkable among rhizobia for having evolved alternative Nod-factor independent nodulation strategies (9). The *Bradyrhizobium* genus has thus been proposed to be the most ancestral symbiont of legumes, from which other rhizobia-legume symbioses evolved (5,10). Importantly, *Bradyrhizobium* bacteria interact symbiotically with legumes of high agronomic importance, among which grain legume, and are the most widely used rhizobia in agriculture. They thus limit the use of nitrogen fertilizers for the promotion of legume crop growth, and more generally, contribute to the nitrogen enrichment of the terrestrial ecosystem, hence playing a very important role in the preservation of the environment.

The *Bradyrhizobium* genus harbouring a single species was created in 1982 to account for their slow growth comparatively to other rhizobia (11) and only three species had been formerly described up until the years 2000. *Bradyrhizobium* has long been considered a ‘taxonomically difficult’ group of organisms due to their highly conserved SSU rRNA gene sequence and to the poor correlation between the groupings formed on the basis of genotypic and phenotypic traits. The emergence of new molecular and bioinformatic methods in the last 20 years as well as the extensive use of whole genome sequence data is now providing a clearer picture of the diversity of this genus. To this date, *Bradyrhizobium* comprises eighty-eight species organized into three major lineages (3) and/or four of the seven supergroups/superclades defined by genome phylogeny (12,13), with intermingled symbiotic and non-symbiotic taxa.

In Australia, *Bradyrhizobium* bacteria are particularly abundant and prevail in Australian acidic and seasonally dry soils (14-18), where they help legumes that constitute about 10% of the estimated 18,000 native plant species, to circumvent nutrient deficiencies and low fertility of Australian soils. Among novel lineages identified in a survey of Australian bradyrhizobia isolated from nodules of indigenous legumes based on SSU rRNA gene sequence analyses (14), *Bradyrhizobium* genospecies L could not unambiguously be affiliated to either of *Bradyrhizobium* major lineages despite its SSU rRNA gene exibiting the *B. elkanii* lineage signature (14).

We here describe a new *Bradyrhizobium* species corresponding to Australian *Bradyrhizobium* genospecies L (14), with BDV5419 for which the complete genome sequence was recently obtained (19) as type strain. *Bradyrhizobium* genospecies L was isolated only once, from *Hardenbergia violacea*. We thus propose naming this novel species *Bradyrhizobium hardenbergiae* sp. nov.

## Isolation and Ecology

Strain BDV5419 was isolated in 1995 from a root nodule of *Hardenbergia violaceae* (Fabaceae: Papilionoideae: Phaeseoleae) collected in Black Mountain Nature Reserve, in Canberra, Australian Capital Territory, Australia (35°16’1.20”S, 149°05’60.00”E), during a survey of rhizobia associated to native shrubby legumes in south-eastern Australia (14). It is a representative of *Bradyrhizobium* genospecies L which was isolated in a single occasion (14). It induces effective nitrogen-fixing nodules on its original host (14).

The strain was short-term cultured on modified yeast extract–mannitol agar (YMA) medium at 25°C (20) and, for long-term preservation, was stored in yeast extract–mannitol broth supplemented with 30 % glycerol (v/v) at −80 °C, as well as lyophilized.

The strain is deposited at the French Collection for Plant-associated Bacteria [Centre International de Ressources Microbiennes (CIRM) – Collection Française des Bactéries associées aux Plantes (CFBP), INRAE] and at the Belgian Coordinated Collections of Microorganisms (BCCM/LMG).

### 16S rRNA gene phylogenies

The first characterization of strain BDV5419 about twenty-five years ago suggested that it represents a novel lineage loosely related to the *B. elkanii* lineage based on SSU rRNA gene sequence phylogeny (14). Many *Bradyrhizobium* species have since been described, which calls for the phylogenetic position of strain BDV5419 within the *Bradyrhizobium* genus to be reassessed. Strain BDV5419 SSU rRNA gene was compared to those of the 88 current *Bradyrhizobium* type species as listed in the List of Prokaryotic names with Standing in Nomenclature (LPSN) database (21) (accessed on March 10^th^ 2024). The SSU rRNA gene sequences were obtained from NCBI Genbank/RefSeq database and analyzed using SeaView version 5.0.4 (22). Sequences were aligned using ClustalO and a maximum likelihood phylogeny was inferred using PhyML version 3.1 (23) with the general time reversible (GTR) model of nucleotide evolution with substitution rates varying over sites according to the invariable sites plus gamma (I + g) distribution as recommended by Abadi *et al*. (24). Internal branch supports were evaluated using the approximate likelihood ratio test based on a Shimodaira-Hasegawa-like procedure (*n*=1,000) (23). The resulting phylogeny confirmed that strain BDV5419 does not group with any formerly described species and represents a novel lineage most basal to the *B. elkanii* lineage (3) that encompasses both the *B. elkanii* and *B. jicamae* supergroups (12, 13) (Figure 1). Including type strains of other *Nitrobacteraceae* type species did not affect the grouping of strain BDV5419 within the *B. elkanii* lineage (Data not shown).

**Figure 1.**
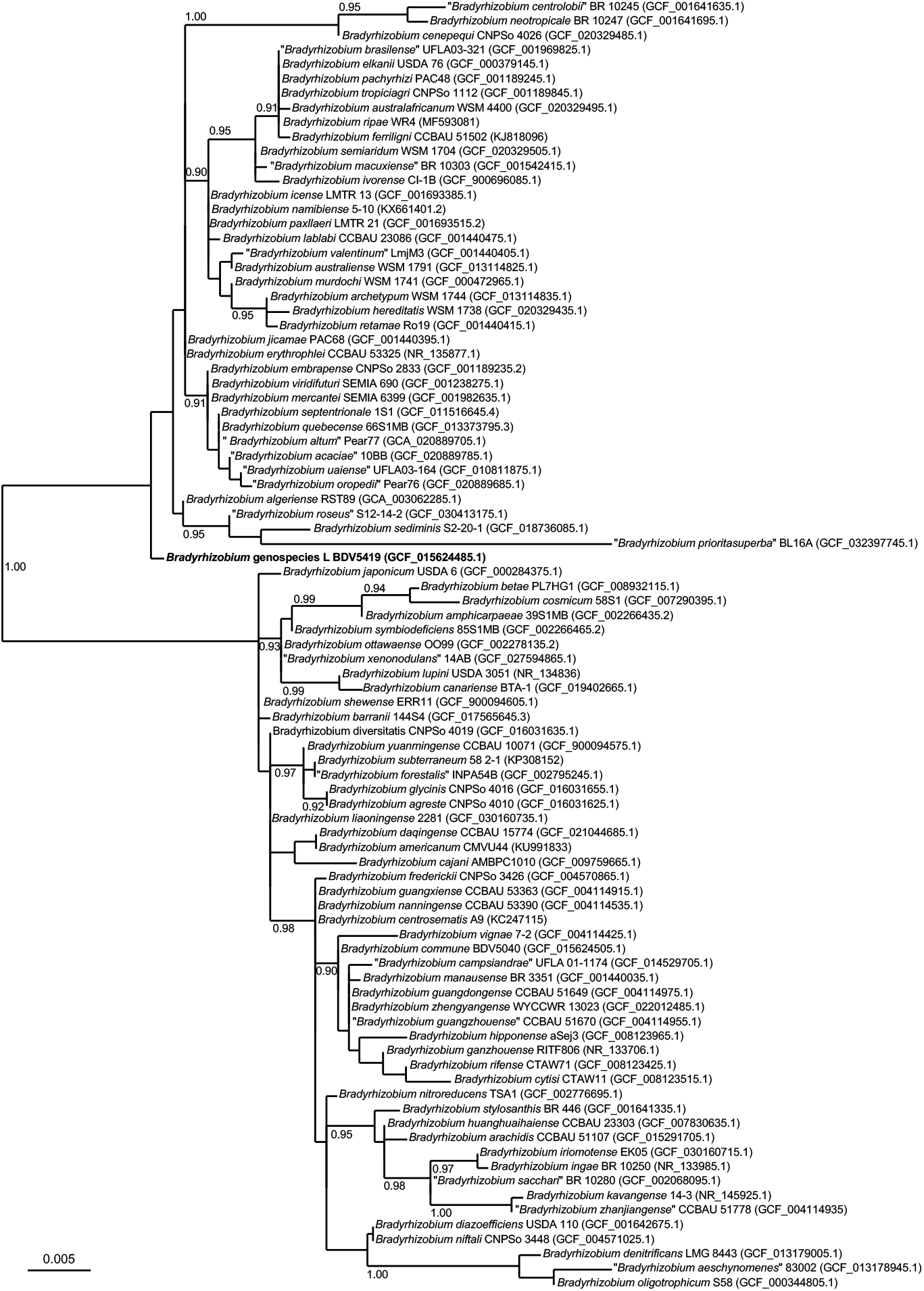
SSU rRNA gene maximum likelihood phylogeny of *Bradyrhizobium* genospecies L BDV5419 among all available *Bradyrhizobium* type strains. The numbers above branches are aLRT support values (only values above 0.90 are shown).

### Genome Features

We further investigated the relationships of strain BDV5419 to known species of *Bradyrhizobium* using whole genome sequence comparison. The complete sequence of strain BDV5419 genome has been reported elsewhere (19). It consists in a 7.40 Mb chromosome with an average G+C content of 64.6 % and encodes 6,879 protein-coding genes, 48 tRNAs, one copy each of the 5S, 16S and 23S rRNA genes, and 94 pseudogenes. Average Nucleotide Identities (ANI) between strain BDV5419 and the 77 *Bradyrhizobium* species with whole genome sequence data (computed using GTDB FastANI Calculator (25) on March 3^rd^ 2024) showed *Bradyrhizobium* genospecies L to be distinct from any known *Bradyrhizobium* species and more closely related to members of the *B. elkanii* supergroup (Table 1). The genome-based taxonomic status of strain BDV5419 was further evaluated using the Type (strain) Genome Server (TYGS) at Leibniz Institute DSMZ (26,27). In connection to the LPSN database, the TYGS determines the closest type strain genomes, calculates intergenomic distances using the Genome BLAST Distance Phylogeny approach (GBDP) and digital DNA-DNA Hybridization (dDDH) values with confidence intervals, and infer a minimum evolution tree with branch support (27). These analyses (performed on March 15^th^ 2024) confirmed that strain BDV5419 constitutes a separate species within the *Bradyrhizobium* genus, most basal with *B. ivorense, B. semiaridum* and “*B. macuxiense*” to the *B. elkanii* supergroup (Figure 2; Table 1).

**Table 1.** Digital DNA-DNA hybridization (dDDH) values with confidence intervals (CI) and Average Nucleotide Identities (ANI) between *Bradyrhizobium* genospecies L BDV5419 and *Bradyrhizobium* type strains with genome data. Genome sequence accession numbers and major characteristics are provided. dDDH values were calculated using TYGS (26, 27). ANIs were computed using GTDB FastANI Calculator (25).

| *Bradyrhizobium*<br>type strain<sup>a</sup> | Genome assembly<br>accession | Genome<br>length (bp) | Genome<br>%G+C | Protein<br>number | dDDH<sup>b</sup><br>(%) | CI (%) | ANI (%) |
| --- | --- | --- | --- | --- | --- | --- | --- |
| "*B. acaciae*" 10BB | GCF_020889785.1 | 8 471 967 | 63,69 | 8102 | 31.70 | [29.3 - 34.2] | 87.68 |
| "*B. aeschynomenes*" 83002 | GCF_013178945.1 | 7 522 254 | 65,42 | 6743 | 23.20 | [20.9 - 25.6] | 82.11 |
| *B. agreste* CNPSo 4010 | GCF_016031625.1 | 7 877 331 | 63,81 | 7292 | 24.00 | [21.7 - 26.5] | 83.10 |
| *B. algeriense* RST89 | GCA_003062285.1 | 8 824 342 | 62,14 | 9374 | 24.90 | [22.6 - 27.4] | 82.92 |
| "*B. altum*" Pear77 | GCA_020889705.1 | 9 758 985 | 63,32 | 9656 | 31.90 | [29.5 - 34.4] | 87.71 |
| *B. amphicarpaceae* 39S1MB | GCF_002266435.2 | 7 044 517 | 64,66 | 6536 | 23.80 | [21.5 - 26.2] | 83.14 |
| *B. arachidis* CCBAU 51107 | GCF_015291705.1 | 9 865 973 | 63,62 | 9313 | 24.60 | [22.3 - 27.1] | 83.56 |
| *B. archetypum* WSM 1744 | GCF_013114835.1 | 7 017 218 | 62,26 | 6656 | 24.10 | [21.8 - 26.6] | 82.73 |
| *B. australafricanum* WSM 4400 | GCF_020329495.1 | 9 684 669 | 63,52 | 9241 | 32.10 | [29.7 - 34.7] | 87.93 |
| *B. australiense* WSM 1791 | GCF_013114825.1 | 7 436 256 | 61,88 | 7200 | 23.90 | [21.6 - 26.3] | 82.56 |
| *B. barranii* 144S4 | GCF_017565645.3 | 11 399 526 | 63,13 | 11285 | 24.10 | [21.8 - 26.5] | 83.05 |
| *B. betae* PL7HG1 | GCF_008932115.1 | 7 419 402 | 64,36 | 7103 | 23.90 | [21.6 - 26.4] | 83.19 |
| "*B. brasiliense*" UFLA03-321 | GCF_001969825.1 | 8 595 048 | 63,93 | 7961 | 32.20 | [29.8 - 34.7] | 87.98 |
| *B. cajani* AMBPC1010 | GCF_009759665.1 | 8 381 257 | 63,96 | 7672 | 24.20 | [21.9 - 26.6] | 83.24 |
| "*B. campsiandrae*" UFLA 01-1174 | GCF_014529705.1 | 9 084 813 | 63,63 | 8764 | 23.70 | [21.4 - 26.2] | 82.93 |
| *B. canariense* BTA-1 | GCF_019402665.1 | 8 561 472 | 63,16 | 8182 | 24.00 | [21.7 - 26.5] | 83.07 |
| *B. cenepequi* CNPSo 4026 | GCF_020329485.1 | 8 472 857 | 62,31 | 8181 | 24.90 | [22.5 - 27.3] | 83.36 |
| "*B. centrolobii*" BR 10245 | GCF_001641635.1 | 10 111 113 | 63,22 | 9545 | 24.00 | [21.7 - 26.5] | 83.10 |
| *B. commune* BDV5040 | GCF_015624505.1 | 7 622 528 | 63,92 | 7107 | 23.80 | [21.5 - 26.2] | 83.15 |
| *B. cosmicum* 58S1 | GCF_007290395.1 | 7 304 136 | 64,33 | 6925 | 23.90 | [21.5 - 26.3] | 83.10 |
| *B. cytisi* CTAW11 | GCF_008123515.1 | 8 854 489 | 63,17 | 8449 | 24.00 | [21.7 - 26.4] | 83.16 |
| *B. daqingense* CCBAU 15774 | GCF_021044685.1 | 7 998 085 | 63,72 | 7566 | 23.80 | [21.5 - 26.2] | 82.82 |
| *B. denitrificans* LMG 8443 | GCF_013179005.1 | 8 333 290 | 64,75 | 7596 | 23.00 | [20.7 - 25.4] | 81.81 |
| *B. diazoefficiens* USDA 110 | GCF_001642675.1 | 9 106 064 | 64,06 | 8493 | 24.20 | [21.9 - 26.7] | 83.38 |
| *B. diversitatis* CNPSo 4019 | GCF_016031635.1 | 8 450 368 | 63,93 | 8050 | 24.20 | [21.9 - 26.7] | 83.04 |
| *B. elkanii* USDA 76 | GCF_000379145.1 | 9 482 467 | 63,72 | 8973 | 32.20 | [29.8 - 34.7] | 87.99 |
| *B. embrapense* CNPSo 2833 | GCF_001189235.2 | 8 267 832 | 63,98 | 7736 | 31.60 | [29.2 - 34.1] | 87.68 |
| "*B. forestalis*" INPA54B | GCF_002795245.1 | 8 252 029 | 63,86 | 7606 | 24.10 | [21.8 - 26.6] | 83.07 |
| *B. frederickii* CNPSo 3426 | GCF_004570865.1 | 8 294 086 | 63,85 | 7853 | 24.00 | [21.7 - 26.5] | 83.26 |
| *B. glycinis* CNPSo 4016 | GCF_016031655.1 | 7 794 411 | 63,73 | 7393 | 24.00 | [21.7 - 26.4] | 82.89 |
| *B. guangdongense* CCBAU 51649 | GCF_004114975.1 | 8 437 991 | 63,26 | 8042 | 24.00 | [21.7 - 26.5] | 82.95 |
| *B. guangxiense* CCBAU 53363 | GCF_004114915.1 | 8 200 121 | 63,91 | 7796 | 23.80 | [21.5 - 26.3] | 83.05 |
| "*B. guangzhouense*" CCBAU 51670 | GCF_004114955.1 | 8 138 177 | 63,39 | 7654 | 23.50 | [21.2 - 25.9] | 82.84 |
| *B. hereditatis* WSM 1738 | GCF_020329435.1 | 7 871 253 | 61,99 | 7462 | 24.10 | [21.8 - 26.6] | 82.54 |
| *B. hipponense* aSej3 | GCF_008123965.1 | 8 826 946 | 62,86 | 8456 | 23.70 | [21.4 - 26.1] | 82.84 |
| *B. huanghuaihaiense* CCBAU 23303 | GCF_007830635.1 | 9 226 856 | 63,94 | 8715 | 24.10 | [21.8 - 26.5] | 83.37 |
| *B. icense* LMTR 13 | GCF_001693385.1 | 8 322 773 | 62,03 | 7965 | 24.20 | [21.9 - 26.7] | 82.84 |
| *B. iriomotense* EK05 | GCF_030160715.1 | 9 789 577 | 62,83 | 9109 | 24.00 | [21.7 - 26.5] | 82.95 |

| ***Bradyrhizobium***<br>**type strain<sup>a</sup>** | **Genome assembly**<br>**accession** | **Genome**<br>**length (bp)** | **Genome**<br>**%G+C** | **Protein**<br>**number** | **dDDH<sup>b</sup>**<br>**(%)** | **CI (%)** | **ANI (%)** |
| --- | --- | --- | --- | --- | --- | --- | --- |
| *B. ivorense* CI-1B | GCF_900696085.1 | 9 412 302 | 64,21 | 8590 | 32.80 | [30.4 - 35.4] | 88.30 |
| *B. japonicum* USDA 6 | GCF_000284375.1 | 9 207 384 | 63,67 | 8701 | 24.10 | [21.8 - 26.5] | 83.12 |
| *B. jicamae* PAC68 | GCF_001440395.1 | 8 738 846 | 62,4 | 8497 | 24.50 | [22.2 - 27.0] | 83.23 |
| *B. lablabi* CCBAU 23086 | GCF_001440475.1 | 8 817 291 | 62,63 | 8553 | 24.50 | [22.1 - 26.9] | 83.13 |
| *B. liaoningense* 2281 | GCF_030160735.1 | 7 899 289 | 63,91 | 8011 | 24.10 | [21.8 - 26.6] | 83.20 |
| "*B. macuxiense*" BR 10303 | GCF_001542415.1 | 8 722 143 | 63,25 | 8202 | 30.90 | [28.5 - 33.4] | 87.16 |
| *B. manausense* BR 3351 | GCF_001440035.1 | 9 145 311 | 62,87 | 8735 | 23.70 | [21.4 - 26.2] | 82.89 |
| *B. mercantei* SEMIA 6399 | GCF_001982635.1 | 8 842 857 | 63,99 | 8142 | 31.80 | [29.4 - 34.3] | 88.10 |
| *B. murdochi* WSM 1741 | GCF_000472965.1 | 7 952 271 | 62,09 | 7570 | 24.10 | [21.8 - 26.6] | 82.74 |
| *B. nanningense* CCBAU 53390 | GCF_004114535.1 | 8 290 088 | 63,44 | 7750 | 23.90 | [21.6 - 26.3] | 83.07 |
| *B. neotropicae* BR 10247 | GCF_001641695.1 | 8 679 329 | 63,6 | 8152 | 24.00 | [21.7 - 26.5] | 83.04 |
| *B. niftali* CNPSO 3448 | GCF_004571025.1 | 9 787 683 | 63,53 | 9214 | 24.20 | [21.9 - 26.7] | 83.38 |
| *B. nitroreducens* TSA1 | GCF_002776695.1 | 8 199 267 | 64,31 | 7569 | 24.20 | [21.9 - 26.7] | 83.37 |
| *B. oligotrophicum* S58 | GCF_000344805.1 | 8 264 165 | 65,13 | 7164 | 23.20 | [20.9 - 25.6] | 82.17 |
| "*B. oropedii*" Pear76 | GCF_020889685.1 | 9 043 029 | 63,57 | 8709 | 31.70 | [29.3 - 34.2] | 87.67 |
| *B. ottawaense* OO99 | GCF_002278135.2 | 8 606 328 | 63,83 | 8012 | 24.20 | [21.8 - 26.6] | 83.25 |
| *B. pachyrhizi* PAC48 | GCF_001189245.1 | 8 711 973 | 63,76 | 8253 | 31.90 | [29.5 - 34.4] | 87.82 |
| *B. paxllaeri* LMTR 21 | GCF_001693515.2 | 8 377 719 | 62,51 | 7901 | 24.40 | [22.1 - 26.9] | 83.28 |
| "*B. prioritassuperba*" BL16A | GCF_032397745.1 | 9 083 454 | 63,05 | 8395 | 21.60 | [19.4 - 24.0] | 80.02 |
| *B. quebecense* 66S1MB | GCF_013373795.3 | 9 172 345 | 63,78 | 8778 | 31.80 | [29.4 - 34.3] | 87.91 |
| *B. retamae* Ro19 | GCF_001440415.1 | 8 466 225 | 61,87 | 8344 | 23.90 | [21.6 - 26.4] | 82.63 |
| *B. rifense* CTAW71 | GCF_008123425.1 | 9 952 932 | 63,02 | 9508 | 23.80 | [21.5 - 26.3] | 83.00 |
| "*B. roseus*" S12-14-2 | GCF_030413175.1 | 7 319 411 | 63,33 | 6905 | 24.20 | [21.9 - 26.6] | 82.91 |
| "*B. sacchari*" BR 10280 | GCF_002068095.1 | 8 698 798 | 63,84 | 8158 | 23.90 | [21.6 - 26.4] | 83.13 |
| *B. sediminis* S2-20-1 | GCF_018736085.1 | 5 554 208 | 63,58 | 5289 | 24.00 | [21.7 - 26.5] | 82.72 |
| *B. semiaridum* WSM 1704 | GCF_020329505.1 | 6 712 655 | 65,1 | 6301 | 32.40 | [29.9 - 34.9] | 88.15 |
| *B. septentrionale* 1S1 | GCF_011516645.4 | 10 410 654 | 63,51 | 9996 | 32.20 | [29.8 - 34.7] | 88.02 |
| *B. shewense* ERR11 | GCF_900094605.1 | 9 163 226 | 63,23 | 8630 | 23.90 | [21.6 - 26.3] | 83.21 |
| *B. stylosanthis* BR 446 | GCF_001641335.1 | 8 801 717 | 64,56 | 8138 | 24.20 | [21.9 - 26.6] | 83.41 |
| *B. symbiodeficiens* 85S1MB | GCF_002266465.2 | 7 039 503 | 64,27 | 6586 | 23.80 | [21.5 - 26.2] | 83.06 |
| *B. tropiciagri* CNPSO 1112 | GCF_001189845.1 | 9 767 314 | 63,49 | 9391 | 31.80 | [29.4 - 34.3] | 87.90 |
| "*B. uaiense*" UFLA03-164 | GCF_010811875.1 | 9 972 761 | 63,26 | 10057 | 31.80 | [29.4 - 34.3] | 87.59 |
| "*B. valentinum*" LmjM3 | GCF_001440405.1 | 8 845 258 | 61,92 | 8740 | 24.40 | [22.1 - 26.9] | 82.90 |
| *B. vignae* 7-2 | GCF_004114425.1 | 8 175 609 | 63,21 | 7762 | 23.90 | [21.6 - 26.4] | 82.90 |
| *B. viridifuturi* SEMIA 690 | GCF_001238275.1 | 8 811 922 | 64,03 | 8844 | 31.60 | [29.2 - 34.1] | 87.78 |
| "*B. xenonodulans*" 14AB | GCF_027594865.1 | 8 276 659 | 63,99 | 7720 | 24.00 | [21.7 - 26.5] | 83.32 |
| *B. yuanmingense* CCBAU 10071 | GCF_900094575.1 | 8 201 522 | 63,77 | 7854 | 23.90 | [21.6 - 26.4] | 82.87 |
| "*B. zhanjiangense*" CCBAU 51778 | GCF_004114935.1 | 9 342 022 | 62,94 | 8858 | 24.00 | [21.7 - 26.4] | 82.98 |
| *B. zhengyangense* WYCCWR 13023 | GCF_022012485.1 | 9 004 494 | 62,87 | 8692 | 23.60 | [21.3 - 26.0] | 82.69 |

<sup>a</sup> Names between quotation marks are not validly published names under the International Code of Nomenclature of Prokaryotes (ICNP)
<sup>b</sup> GBDP formula d4 corresponding to the sum of all identities found in High Scoring segment Pairs (HSPs) divided by overall HSP length (29)

**Figure 2.**
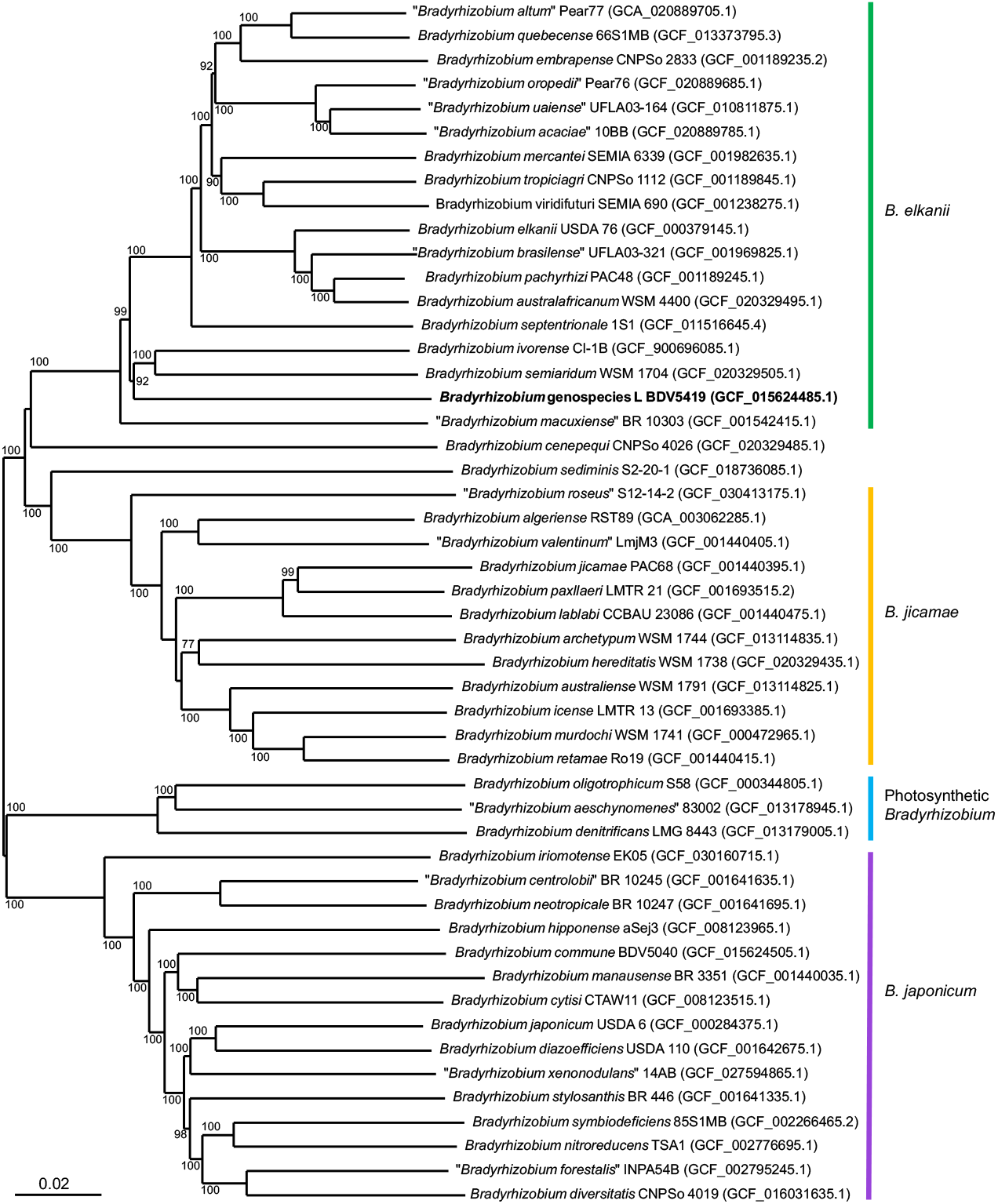
Genome phylogeny of *Bradyrhizobium* genospecies L BDV5419 with all available type strains of *B. elkanii, B. jicamae* and Photosynthetic *Bradyrhizobium* supergroups and representatives of the species diversity of the *B. japonicum* supergroup. Supergroup colouring is according to Avontuur *et al*. (12). The tree was inferred with FastME from GBDP distances calculated from genome sequences. The branch lengths are scaled in terms of GBDP distance formula d5 (derived from d4 distances). The numbers above branches are GBDP pseudo-bootstrap support values from 100 replications.

The initial characterization of *Bradyrhizobium* genospecies L relied on SSU rRNA gene restriction fragment length polymorphisms and sequencing followed by SSU rDNA phylogeny to assess its relationships to then known rhizobial lineages (14). *Bradyrhizobium* genospecies L appeared to be related to, although distant from, *B. elkanii*. The reevaluation of the SSU rRNA gene phylogenetic position of strain BDV5419 among an up-to-date sampling of *Bradyrhizobium* species confirms that it represents a distinct *Bradyrhizobium* species, more closely related to the *B. elkanii* lineage (3) than to either the *B japonicum* or the Photosynthetic *Bradyrhizobium* lineages. However, the SSU rRNA gene phylogeny does not resolve further the relationships of *Bradyrhizobium* genospecies L with members of the *B. elkanii* and *B. jicamae* supergroups that were later individualized using whole genome phylogenetic analyses (12, 13). Overall genomic relatedness indices (OGRI) dDDH and ANI between strain BDV5419 and the 77 *Bradyrhizobium* type strains for which a genome sequence is currently available confirmed the clear delineation of strain BDV5419, all pairwise comparisons being well below the inter-species 70% (dDDH) and 95-96% (ANI) threshold values (28). In contrast to the SSU rRNA gene sequence data, highest OGRI values unambiguously identified the species of the *B. elkanii* supergroup as the closest relatives of BDV5419 (dDDH values ranging from 30.9 to 32.8% and ANI values from 87.16 to 88.3%). Indeed, OGRIs between strain BDV5419 and species of the *B. jicamae* supergroups (23.9-24.9% for dDDH and 82.54-83.28% for ANI) were in the similar, intermediary, value range than that observed with species of the *B. japonicum* supergroup (23.5-24.6% for dDDH and 82.69-83.56% for ANI). Strain BDV5419 was more highly divergent from species of the Photosynthetic supergroup (23.0-23.2% for dDDH and 81.81-82.17% for ANI), and most from *B. prioritasuperba*. Interestingly, OGRIs for this species were in the range of dDDH and ANI values observed between strain BDV5419 and type species of other *Nitrobacteraceae* (Table 2).

**Table 2.** Digital DNA-DNA hybridization (dDDH) values with confidence intervals (CI) and Average Nucleotide Identities (ANI) between *Bradyrhizobium* genospecies L BDV5419 and type strains of *Nitrobacteraceae* type species with genome data. Genome sequence accession numbers and major characteristics are provided. dDDH values were calculated using TYGS (26, 27). ANIs were computed using GTDB FastANI Calculator (25).

| *Nitrobacteraceae* type species<br>type strain | Genome assembly<br>accession | Genome<br>length (bp) | Genome<br>%G+C | Protein<br>number | dDDH<sup>a</sup><br>(%) | CI (%) | ANI (%) |
| --- | --- | --- | --- | --- | --- | --- | --- |
| *Afipia felis* B-91-007352 | GCF_900445155.1 | 4 195 257 | 60.72 | 4049 | 20.90 | [18.7 - 23.4] | 79.47 |
| *Nitrobacter winogradskyi* Nb-255 | GCF_000012725.1 | 3 402 093 | 62.05 | 3262 | 22.10 | [19.8 - 24.5] | 79.61 |
| *Pseudolabrys taiwanensis* CC-BB4 | GCF_003367395.1 | 5 593 741 | 64.39 | 5286 | 19.90 | [17.7 - 22.3] | 78.37 |
| *Pseudorhodoplanes sinuspersici* RIPI 110 | GCF_002119765.1 | 5 971 615 | 60.32 | 5795 | 19.60 | [17.4 - 22] | 77.79 |
| *Rhodoplanes roseus* 941 | GCF_003258865.1 | 6 879 536 | 69.35 | 6828 | 19.90 | [17.7 - 22.3] | 78.00 |
| *Rhodopseudomonas palustris* 2.1.6 | GCF_034479375.1 | 5 305 909 | 65.98 | 4736 | 22.60 | [20.3 - 25.1] | 81.67 |
| *Tardiphaga robiniae* LMG 26467 | GCF_013359755.1 | 6 318 076 | 61.51 | 5935 | 21.40 | [19.2 - 23.8] | 80.30 |
| *Undibacter mobilis* GY_H | GCF_003367195.1 | 4 098 220 | 63.28 | 3910 | 19.90 | [17.7 - 22.3] | 78.30 |
| *Variibacter gotjawalensis* GJW-30 | GCF_002355335.1 | 4 586 237 | 62.23 | 4441 | 19.90 | [17.7 - 22.3] | 77.51 |

<sup>a</sup> GBDP formula d4 corresponding to the sum of all identities found in High Scoring segment Pairs (HSPs) divided by overall HSP length (29)

Among species of the *B. elkanii* supergroup, closest relatives to strain BDV5419 were *B. ivorense, B. semiaridum, B. septentrionale, B. brasilense* and *B. elkanii* according to the dDDH values and *B. ivorense, B. semiaridum, B. mercantei*, and *B. septentrionale* according to the ANI values. Accordingly, the genome-based phylogeny identifies strain BDV5419 as a novel species of the *B. elkanii* supergroup. In turn, alike the SSU rRNA gene phylogeny, it places it among the basal and most divergent lineages of this group with strain BDV5419 being most closely related to *B. ivorense* and *B. semiaridum* although separated from them by long branches.

SSU rRNA gene phylogeny and OGRIs whilst providing valuable indication of bacterial inter-species relatedness failed to establish unambiguously the position of strain BDV5419 within the *Bradyrhizobium* genus. The complex evolutionary signal linking strain BDV5419 to the other *Bradyrhizobium* species could only be recovered using whole genome sequence-based phylogeny. This supports the recent proposition to center the taxonomic description of bacterial taxa on genome phylogeny using whole genome sequence data (28).

### Physiology and Chemotaxonomy

Phenotypic tests were performed at 28 °C and included ability to grow in modified YMA medium (20) with the addition of 0.0%, 0.5%, 1 %, 1.5% or 2% NaCl, in modified YMA medium at pH 4.0, 5.0, 6.0, 6.5, 6.8, 7.0, 7.5, 8.0, 8.5 or 9.0, and in solid Luria–Bertani (LB) medium. The ability to grow at various temperature (15°C, 20°C, 28°C, 34°C, 37°C and 40°C) was also tested. All analyses were conducted in triplicate and evaluated over the course of 10 days of incubation. Growth of strain BDV5419 is observed at 15-34°C, pH from 6.5-8.5 and only NaCl concentration below 0.5 %. Optimal growth was obtained at 28°C, pH 6.5, and in the absence of NaCl. Antibiotic tolerance was evaluated using the disc-diffusion method on YMA plates with the following antibiotics (concentration per disc): ampicillin (10 μg.ml^-1^), chloramphenicol (30 μg.ml^-1^), erythromycin (15 μg.ml^-1^), gentamycin (30 μg.ml^-1^), kanamycin (30 μg.ml^-1^), nalidixic acid (30 μg.ml^-1^), rifampicine (10 μg.ml^-1^), and streptomycin (10 μg.ml^-1^). Strain BDV5419 could tolerate kanamycin, rifampicine and streptomycin. Enzymatic activities and carbon (C) source utilization capacities were evaluated using the API 20NE kit (BioMérieux) following the manufacturer’s recommendations. YM without mannitol was used as basal medium and bromothymol blue as indicator of acid or alkaline reaction when the use of each C source was evaluated. The main properties are included in the species description.

### Protologue

*Bradyrhizobium hardenbergiae* ((har.den.ber’gi.ae. N.L. gen. n. adj. *hardenbergiae*, of *Hardenbergia violaceae*, from which the type strain was isolated).

Cells are Gram-negative, non-spore-forming aerobic rods. Colonies are circular, mucoid, opaque white and convex on YMA medium after 7 days of growth at 28 °C. Optimum growth occurs at pH 6.5 and 28 °C, although growth will also occur at pH 6.0-8.5, and 15-34°C. Strain BDV5419 does not grow in the presence of NaCl or in LB medium. It is resistant to kanamycin (30 μg/ml), rifampicine (10 μg/ml) and streptomycin (10 μg/ml). The type strain is susceptible to ampicillin (10 μg/ml), chloramphenicol (30 μg/ml), erythromycin (15 μg/ml), gentamycin (30 μg/ml) and nalidixic acid (30 μg/ml). Strain BDV5419 tests negative for indole production, glucose fermentation and gelatin hydrolysis. The type strain tests positive for nitrate reduction, urease, arginine dihydrolase, β-galactosidase and β-glucosidase activities. This strain is capable of assimilating D-glucose, D-maltose, D-mannose, L-arabinose, malic acid, phenylacetic acid and trisodium citrate as carbon source as carbon source but tested negative for assimilation of D-mannitol, N-acetylglucosamine, adipic acid, capric acid and potassium gluconate is negative. It may induce effective nitrogen-fixing nodules on its original host.

The type strain is BDV5419^T^ (=CFBP 9111^T^ = LMG 32897^T^), isolated from nodules of *Hardenbergia violacea* in Australia. The DNA G+C content of strain BDV5419^T^ is 64.6 mol%. Molecular accession numbers for type strain are: SSU rDNA gene (Z94821), chromosome (CP061378), genome assembly (GCA_015624485).

## Abbreviations

ACT: Australian Capital Territory
ANI: Average Nucleotide Identity
ICNP: International Code of Nomenclature of Prokaryotes
dDDH: digital DNA-DNA Hybridization
GBDP: Genome BLAST Distance Phylogeny
LB: Luria-Bertani
LPSN: List of Prokaryotic names with Standing in Nomenclature
OGRI: Overall Genome Relatedness Index
YMA: yeast extract–mannitol agar

## Notes

### Competing Interest Statement

The authors have declared no competing interest.

## References

1. Delgado-Baquerizo M, Oliverio AM, Brewer TE, Benavent-González A, Eldridge DJ, BardgeÇ RD, et al. A global atlas of the dominant bacteria found in soil. Science. 2018;359(6373):320–5. DOI: 10.1126/science.aap9516

2. VanInsberghe D, Maas KR, Cardenas E, Strachan CR, Hallam SJ, Mohn WW. Non-symbiotic Bradyrhizobium ecotypes dominate North American forest soils. ISME J. 2015;9(11):2435–41. DOI: 10.1038/ismej.2015.54

3. Lindström K, Aserse AA, Mousavi AS. Evolution and taxonomy of nitrogen-fixing organisms with emphasis on rhizobia. In: de Bruijn FJ, editor. Biological Nitrogen Fixation. John Wiley & Sons Ltd; 2015. p. 21–38. DOI: 10.1002/9781119053095.ch3

4. Sprent JI, Ardley J, James EK. Biogeography of nodulated legumes and their nitrogen-fixing symbionts. New Phytol. 2017;215(1):40–56. DOI: 10.1111/nph.14474

5. Parker MA. The spread of Bradyrhizobium lineages across host legume clades: from Abarema to Zygia. Microb Ecol. 2015;69:630–40. DOI: 10.1007/s00248-014-0503-5

6. Andrews M, Andrews ME. Specificity in Legume-Rhizobia Symbioses. Int J Mol Sci. 2017;18(4):705. DOI: 10.3390/ijms18040705

7. Trinick MJ. Symbiosis between Rhizobium and the non-legume, Trema aspera. Nature. 1973;244(5416):459–60. DOI: 10.1038/244459a0

8. Trinick MJ, Hadobas PA. Biology of the Parasponia-Bradyrhizobium symbiosis. Plant Soil. 1988;110(2):177–85. DOI: 10.1007/BF02226797

9. Okazaki S, TiÇabutr P, Teulet A, Thouin J, Fardoux J, Chaintreuil C, et al. Rhizobium-legume symbiosis in the absence of Nod factors: two possible scenarios with or without the T3SS. ISME J. 2016;10(1):64–74. DOI: 10.1038/ismej.2015.103

10. Hungria M, Menna P, Marçon Delamuta JR. Bradyrhizobium, the ancestor of all rhizobia: phylogeny of housekeeping and nitrogen-fixation genes. In: Bruijn FJd, editor. Biological Nitrogen Fixation: John Wiley & Sons, Inc; 2015. p. 191–202. DOI: 10.1002/9781119053095.ch18

11. Jordan DC. Transfer of Rhizobium japonicum Buchanan 1980 to Bradyrhizobium gen. nov., a genus of slow-growing, root nodule bacteria from leguminous plants. Int J Syst Bacteriol. 1982;32:136–9. DOI: 10.1099/00207713-32-1-136

12. Avontuur JR, Palmer M, Beukes CW, Chan WY, Coetzee MPA, Blom J, et al. Genome-informed Bradyrhizobium taxonomy: where to from here? Syst Appl Microbiol. 2019;42(4):427–39.

13. Ormeño-Orrillo E, Marànez-Romero E. A Genomotaxonomy View of the Bradyrhizobium Genus. Front Microbiol. 2019;10:1334. DOI: 10.3389/fmicb.2019.01334

14. Lafay B, Burdon JJ. Molecular diversity of rhizobia occurring on native shrubby legumes in Southeastern Australia. Appl Environ Microbiol. 1998;64(10):3989–97. DOI: 10.1128/AEM.64.10.3989-3997.1998

15. Lafay B, Burdon JJ. Small-subunit rRNA genotyping of rhizobia nodulating Australian Acacia spp. Appl Environ Microbiol. 2001;67(1):396–402. DOI: 10.1128/AEM.67.1.396-402.2001

16. Lafay B, Burdon JJ. Molecular diversity of rhizobia nodulating the invasive legume CyCsus scoparius in Australia. J Appl Microbiol. 2006;100:1228–38. DOI: 10.1111/j.1365-2672.2006.02902.x

17. Lafay B, Burdon JJ. Molecular diversity of legume root-nodule bacteria in Kakadu National Park, Northern Territory, Australia. PLoS One. 2007;2(3):e277. DOI: 10.1371/journal.pone.0000277

18. Stępkowski T, Watkin E, McInnes A, Gurda D, Gracz J, Steenkamp ET. Distinct Bradyrhizobium communities nodulate legumes native to temperate and tropical monsoon Australia. Mol Phylogenet Evol. 2012;63(2):265–77. DOI: 10.1016/j.ympev.2011.12.020

19. Oger-Desfeux C, Briolay J, Oger PM, Lafay B. Complete genome sequence of Bradyrhizobium sp. strain BDV5419, representative of Australian genospecies L. Microbiol Resour Announc. 2021;10(8):e00042–21.

20. Vincent JM. A manual for the practical study of root-nodule bacteria. International biological programme handbook No. 15. Oxford, England: Blackwell Science Publications; 1970.

21. Parte AC, Sarda Carbasse J, Meier-Kolthoff JP, Reimer LC, Göker M. List of Prokaryotic names with Standing in Nomenclature (LPSN) moves to the DSMZ. Int J Syst Evol Microbiol. 2020;70(11):5607–12. DOI: 10.1099/ijsem.0.004332

22. Gouy M, Tannier E, Comte N, Parsons DP. Seaview Version 5: A multiplatform software for multiple sequence alignment, molecular phylogenetic analyses, and tree reconciliation. Methods Mol Biol. 2021;2231:241–60. DOI: 10.1007/978-1-0716-1036-7_15

23. Guindon S, Dufayard J-F, Lefort V, Anisimova M, Hordijk W, Gascuel O. New algorithms and methods to estimate maximum-likelihood phylogenies: assessing the performance of PhyML 3.0. Syst Biol. 2010;59(3):307–21. DOI: 10.1093/sysbio/syq010

24. Abadi S, Azouri D, Pupko T, Mayrose I. Model selection may not be a mandatory step for phylogeny reconstruction. Nat Commun. 2019;10(1):934. DOI: 10.1038/s41467-019-08822-w

25. Jain C, Rodriguez-R LM, Phillippy AM, Konstantinidis KT, Aluru S. High throughput ANI analysis of 90K prokaryotic genomes reveals clear species boundaries. Nat Commun. 2018;9:5114. DOI: 10.1038/s41467-018-07641-9

26. Meier-Kolthoff JP, Göker M. TYGS is an automated high-throughput platform for state-of-the-art genome-based taxonomy. Nat Commun. 2019;10(1):2182. DOI: 10.1038/s41467-019-10210-3

27. Meier-Kolthoff JP, Carbasse JS, Peinado-Olarte RL, Göker M. TYGS and LPSN: a database tandem for fast and reliable genome-based classification and nomenclature of prokaryotes. Nucleic Acids Res. 2022;50(D1):D801–D7. DOI: 10.1093/nar/gkab902

28. Riesco R, Trujillo ME. Update on the proposed minimal standards for the use of genome data for the taxonomy of prokaryotes. Int J Syst Evol Microbiol. 2024;74(3):006300. DOI: 10.1099/ijsem.0.006300

29. Meier-Kolthoff JP, Auch AF, Klenk HP, Göker M. Genome sequence-based species delimitation with confidence intervals and improved distance functions. BMC Bioinformatics. 2013;14:60. DOI: 10.1186/1471-2105-14-60

